# Individualized Electrode Subset Improves the Calibration Accuracy of an EEG P300-design Brain-Computer Interface for People with Severe Cerebral Palsy

**DOI:** 10.1101/2023.03.22.533775

**Authors:** Si Long Jenny Tou, Seth A. Warschausky, Petra Karlsson, Jane E. Huggins

## Abstract

**Objective:** This study examined the effect of individualized electroencephalogram (EEG) electrode location selection for non-invasive P300-design brain-computer interfaces (BCIs) in people with varying severity of cerebral palsy (CP).

**Approach:** A forward selection algorithm was used to select the best performing 8 electrodes (of an available 32) to construct an individualized electrode subset for each participant. BCI accuracy of the individualized subset was compared to accuracy of a widely used default subset.

**Main Results:** Electrode selection significantly improved BCI calibration accuracy for the group with severe CP. Significant group effect was not found for the group of typically developing controls and the group with mild CP. However, several individuals with mild CP showed improved performance. Using the individualized electrode subsets, there was no significant difference in accuracy between calibration and evaluation data in the mild CP group, but there was a reduction in accuracy from calibration to evaluation in controls.

**Significance:** The findings suggested that electrode selection can accommodate developmental neurological impairments in people with severe CP, while the default electrode locations are sufficient for many people with milder impairments from CP and typically developing individuals.

## 1. Background

P300-design electroencephalogram (EEG) brain-computer interface (BCI) interprets brain activity to translate users’ intentions into computer commands to type or generate speech for communication without physical movement [1]. The P300 is a positive in magnitude component occurring about 300 ms after perception of an expected target stimulus that occurs infrequently and randomly. However, the P300 BCI design does not specifically check for P300s. Instead it will detect any event-related potential (ERP) components including the P300, N200, etc. [2]. The EEG P300-design BCI was first proposed in Farwell and Donchin, 1988 with only one electrode location used [1]. Subsequent studies using additional electrodes showed improved BCI performance [3, 4], and recent P300-design BCIs usually collect data from a set of multiple electrodes with fixed default locations. Some of the latest P300-design BCI studies use up to 64 electrodes [4,5]. A larger electrode set, however, increases cost and setup time, and makes the BCI less practical for end users [6–8]. In addition, increasing the size of an electrode subset may only yield minimal improvement in accuracy beyond a subset of size 4 for typically developing individuals [9].

Studies have shown that P300-design BCIs perform better when used by typically developing participants compared to those with neurological conditions [10,11]. The mechanisms underlying these differences have not been clearly identified. However, this disparity may stem in part from the effects of underlying physiological differences in brain function. This raises concerns that commercialization efforts that seek to minimize the number of electrodes will result in BCIs that do not work for people with physical impairments. If electrode selection can create individualized electrode subsets that provide better signal detection, while using a minimal number of electrodes, BCI can become more accurate, lower in cost, and faster to set up. Using an effective electrode selection algorithm to obtain custom electrode subsets has the potential to remove barriers to BCI use outside the laboratory environment.

Various electrode selection methods are discussed in the literature on EEG-BCI. Alotaiby et al. [12] summarized popular filtering, wrapping, embedded, and hybrid electrode selection techniques for EEG. Filtering techniques use independent criteria to assess the usefulness of electrodes. Wrapper techniques, on the other hand, depend on a classification algorithm or machine learning to evaluate an individual’s electrode subset. While filtering techniques are computationally much cheaper and faster than wrapper techniques, its limitation is often its low accuracy [12]. Examples of filtering techniques discussed in Alotaiby et al. include minimizing within-class variance, maximizing between-class variance, maximizing entropy, and using common spatial pattern filter coefficients. Instances of wrapper techniques include Support Vector Machine (SVM) to rank the contributions of each selected electrode, and Regularized Linear Discriminant Classifier (RLDC) [12]. Embedded techniques incoporate electrode selection into classifier construction. Hybrid is a combination of the above techniques [12]. Other methods include using Principal Component Analysis (PCA) to assess contributions of each electrode [13], inconsistencies of classifiers [14], and more recently backtracking search optimization [15]. Some electrode selection methods that have been evaluated with P300-design BCI data include exhaustive search [6], forward selection [6,9], backward elimination [6,9], backward elimination based on ‘‘signal to signal-plus-noise ratio” [8], n-forward-m-reverse search [6], jumpwise selection [6], phase measurement [16], Bayesian linear discriminant analysis with a sparse prior [7], artificial neural networks [17], and genetic algorithms [17–19].

This study examines the benefits to P300-BCI performance of individualizing electrode locations with the aim to better match potentially atypical neuroanatomy and neurophysiology of individual participants with cerebral palsy (CP). Specifically, this study is an offline investigation of the effect of a custom individualized electrode subset on P300-BCI accuracy for participants with severe CP, mild CP, and typically developing controls. We hypothesize that people with CP can benefit from electrode selection by 1) improvement in BCI accuracy; and 2) the ability to use a smaller electrode subset size without jeopardizing performance. To our knowledge there is no systematic analysis of P300-design BCI accuracy using electrode selection for populations with CP in the existing literature.

## 2. Methods

### 2.1. Data for Analysis

We conducted offline analysis on data from two different protocols of BCI use to access a 4-choice vocabulary test. Data in Protocol 1 was collected from participants with mild CP and typically developing controls, described in Alcaide et al. [20] and Warchausky et al. [21]. Protocol 2 involved participants with severe CP, described in [22]. In particular, participants in Protocol 1 were required to have sufficient physical abilities to take the standard, paper version of a forced-choice vocabulary test. This meant that they were required to either be able to say the number of the desired answer or to reliably point to the answer. Participants in Protocol 2 were required to be unable to access the standard version of the test.

In both protocols, data was collected from a 32 gel electrode EEG cap (F3, Fz, F4, FC5, FC3, FC1, FCz, FC2, FC4, FC6, T7, C5, C3, C1, Cz, C2, C4, C6, T8, CP5, CP3, CP1, CPz, CP2, CP4, CP6, P3, Pz, P4, PO7, PO8, Oz), with only 16 electrodes (default 16 in Figure 5) used for in-session calibration (F3, Fz, F4, T7, C3, Cz, C4, T8, CP3, CP4, P3, Pz, P4, PO7, PO8, Oz). EEG was recorded with two synchronized g.USBamp EEG amplifiers (Guger Technologies, Austria).

The layout of the interface was chosen to mimic the layout of the paper PPVT-IV [27] except that the numeric labels were moved from under each picture to the corner of each picture to provide maximum separation between stimuli. Likewise, the cancel button was position at the center of the screen to provide maximum separation from all picture labels.

Each question (trial) of the vocabulary testing format included the audio presentation of a recorded word (audio prompt) and the presentation on-screen of four pictures. Literacy was not an inclusion/exclusion criteria and no words were presented in written form. Words for calibration were selected to be easy to match to the pictures. During recording of the calibration data, the user’s attention was drawn to the picture that matched the audible word by leaving that picture in color and changing the other pictures to grayscale. Each picture had a labeled corner in which gray text on a black background intensified (flashed) to bright white as the stimulus to elicit the P300. The labels flashed individually in a random order. Each label flashed once before any label repeated. The EEG after each flash was analyzed for the presence or absence of a P300 response (see Fig. Appendix A.1). Each label is flashed 10 times for each trial.

#### 2.1.1. Differences between Protocols

The P300 stimuli for Protocol 1 had an additional continuously flickering checkerboard border (Figure 2, left) that was intended to elicit a steady-state visual evoked potential (SSVEP). Offline SSVEP analysis of the Protocol 1 data did not show significant SSVEP responses and incorrect monitor refresh rates during data collection were suspected. This border was not used in Protocol 2 (Figure 2, right). Stimulus durations were different as a result of the different sample rates but were matched as closely as possible (duty cycle of 166.67ms for Protocol 1 vs 156.25ms for Protocol 2). P300 stimulus flashes for Protocol 1 were 50ms in length with 116.7ms between flashes. P300 stimulus flashes for Protocol 2 were 62.5ms in length with 93.75ms between flashes. The duty cycle for the stimuli was chosen to match that in our past BCI studies [24]. The duration of the flashes was increased to accommodate potentially slower visual processing speed for people with CP [28].

**Figure 1:**
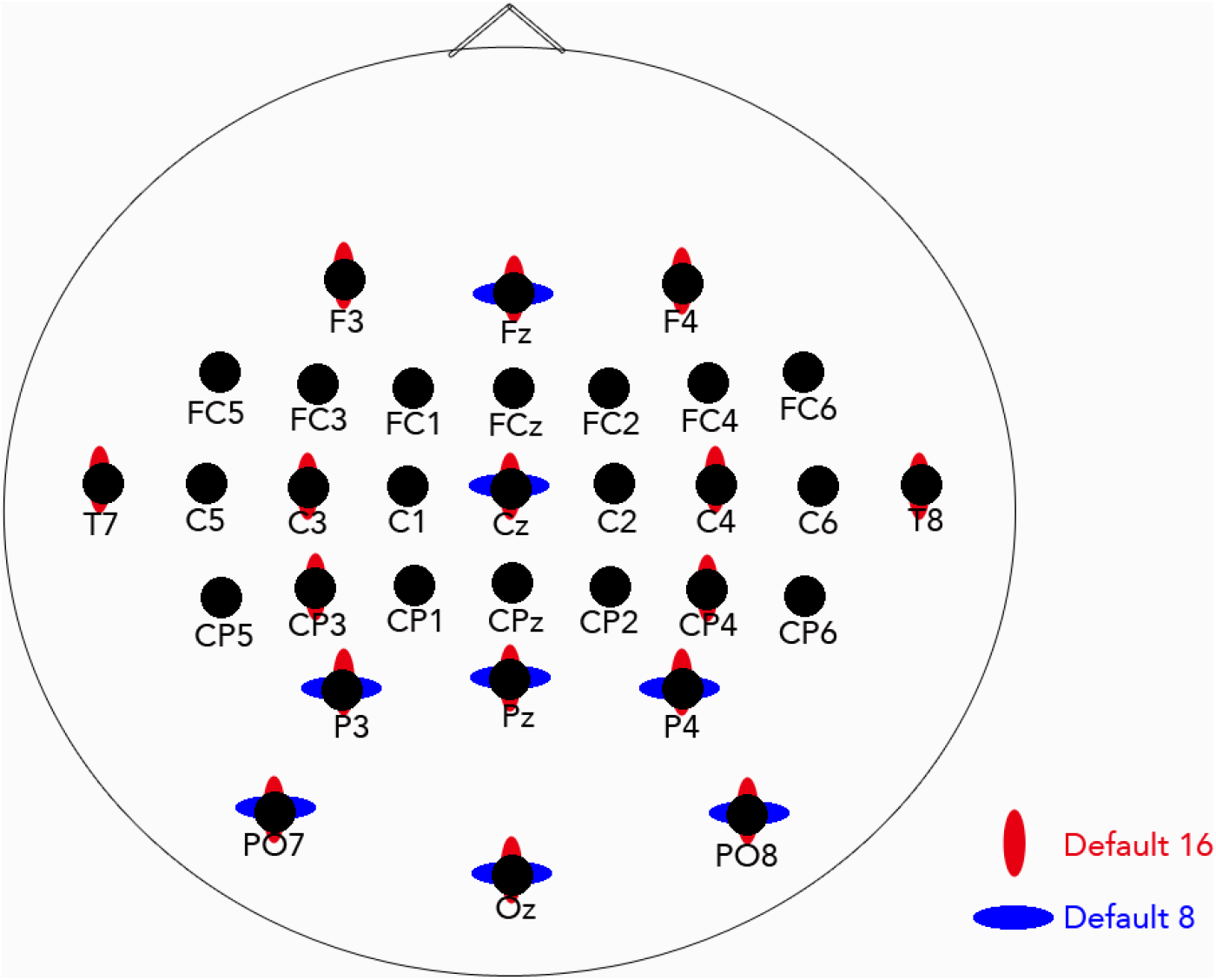
Locations of the 32 electrodes with the default 16 [9, 23–25] and default 8 [26] electrodes marked.

**Figure 2:**
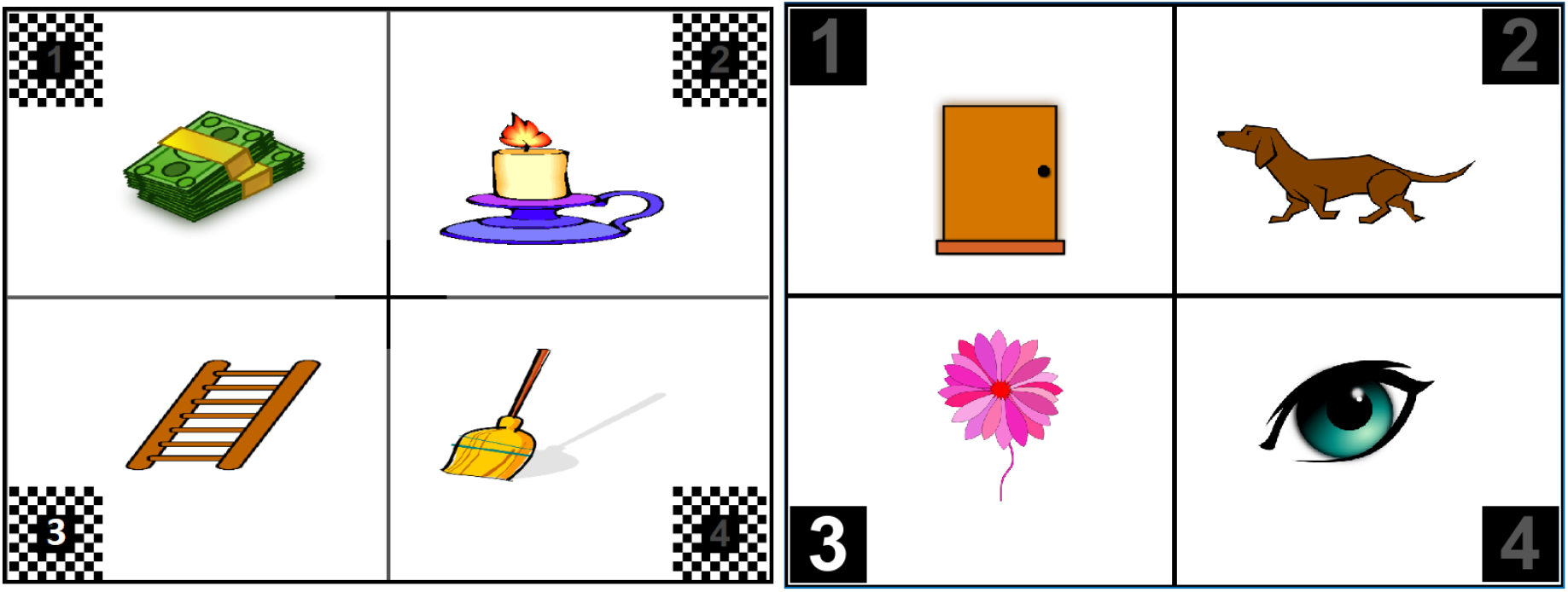
Screen layout for the BCI-presented vocabulary test format for Left: Protocol 1; Right: Protocol 2. Since PPVT-IV [27] images are copy-righted, images shown are those used in calibration.

For protocol 1, the training vocabulary pictures were selected to mimic the PPVT, but to not include words in the actual PPVT and to be relatively simple illustrations of words that we expected participants to recognize. For protocol 2, pictures were chosen in coordination with our collaborators in Australia to select words and illustrations that would be familiar to participants in both countries [22].

Inclusion criteria for Protocol 1 were ages 8-29 years. The original recruitment age was originally restricted to 10-29 years, but the upper bound was later relaxed to 45 years. The original recruitment criteria for Protocol 2 was originally restricted to 10-29 years, but later relaxed to 45 years.

Audio prompts for Protocol 1 had only the word to find. Audio prompts for Protocol two also included the instruction “Find” as well as the word. The words and images used for the protocols were different since Protocol 2 words were carefully selected for awareness by those with severe impairments.

In Protocol 1, EEG was sampled at 600 Hz and calibration data consisted of at least 60 trials in separate files of 30 trials each. In Protocol 2, EEG was sampled at 256 Hz and calibration data was intended to include 30 trials. These trials were separated into shorter trial blocks intended to accommodate limited attention spans.

### 2.2. Participants in Protocol 1

Data were obtained from 17 participants with CP (5 females and 12 males, age 14.7±5.6 years) and 10 typically developing participants (6 females and 4 males, age 14.7±4.3 years) [20,21]. The Institutional Review Board of the University of Michigan approved the protocol (HUM00012968). Informed consent was obtained from the human participants, a parent, or guardian. Inclusion criteria for both the group with CP and the typically developing group were sufficient vision and sufficient speech or movement to participate in the standardized version of the Peabody Picture Vocabulary Test Fourth Edition (PPVT-IV) [27]. Exclusion criteria included 1) history of moderate or severe acquired brain injury or other major neurological condition such as stroke, encephalitis, or refractory seizure disorder (for those with CP, this refers to events subsequent to the onset and diagnosis of CP), 2) major psychiatric disorder such as major depression, severe anxiety or psychosis that precluded participation, and 3) for those under age 18, inability of the parent/guardian to complete a child history. In the group with CP, one participant was taking baclofen and one was taking sertraline. In the typically developing group, one participant was taking sertraline.

In the group with CP, primary tone in all participants was spasticity, 60.0% had hemiplegia and 40% diplegia. Functional classifications were obtained with the Gross Motor Functional Classification System (GMFCS) [29], the Manual Ability Classification System (MACS) [30], and the Oral Communication Communication Function Classification System (CFCS) [31]. Score distributions are described in table 1.

**Table 1:** Distributions of participants by functional classification sytem in Protocol 1 and Protocol 2. For participants who reported more than one levels for a classification system, they are categorized into the less severe level.

|  |  | Level I | Level II | Level III | Level IV | Level V | Did not report |
| --- | --- | --- | --- | --- | --- | --- | --- |
| Protocol 1<br>(Mild CP) | GMFCS | 64.7% | - | 11.8% | 5.9% | 5.9% | 11.8% |
|  | MACS | 23.5% | 47.1% | 11.8% | 11.8% | - | 5.9% |
|  | CFCS | 70.6% | 11.8% | 5.9% | 5.9% | - | 5.9% |
| Protocol 2<br>(Severe CP) | GMFCS | - | - | - | 16.7% | 83.3% | - |
|  | MACS | - | - | - | 8.3% | 91.7% | - |
|  | CFCS | - | 4.2% | - | 45.8% | 50% | - |

Among the participants, 10 with CP (2 females and 8 males, age 17.4±5.5 years) and all typically developing controls completed the study. However, for seven participants with CP (3 females and 4 males, age 10.9±3.0 years), the BCI could not be calibrated at performance threshold of 75% with the study settings, even with multiple repetitions of the calibration phase. It is noteworthy that those for whom a calibration could not be found included many of the younger participants in the group. In addition, one participant with CP who was able to use the BCI only had 30 trials of calibration data. All participants in Protocol 1 are included in the electrode selection calibration analysis as detailed in section 2.6; only participants who had a successful calibration phase (9 participants with CP, 10 participants not affected by CP) then completed the testing phase as detailed in section 2.7.

### 2.3. Participants in Protocol 2

In Protocol 2, usable BCI calibration data from 24 participants with severe CP (16 females and 8 males, age 18.3±8.4 years) were obtained. Participants were part of a study comparing access to a multiple-choice test with BCI-access and eye-gaze-interface-access [22]. The Institutional Review Board of the University of Michigan approved the protocol (HUM00130187) as did the Cerebral Palsy Alliance Human Research Ethics Committee (approval number: 2017_05_02). Informed consent was obtained from the human participants, a parent, or guardian. The participants for this study were required to be unable to access the paper version of the multiple-choice test.

Sixteen participants were from the USA and 8 from Australia. Primary tone for 17 participants (70.8%) was spasticity, four participants (16.7%) was dystonia, and three participants (12.5%) had diagnoses of both spasticity and dystonia. Participant characteristics are described in table 1.

Among the 24 participants, there were five for whom calibration with the 16 electrodes used for the online study met the performance threshold of 75% accuracy for successful calibration.

### 2.4. Offline Data Preprocessing

All data were processed with a modified BCI2000 (version 3) P300 Classifier tool (bci2000.org) [32]. All functionality and the implementation of step-wise linear discriminant analysis (SWLDA) classification from the original BCI2000 P300 Classifier were preserved, with the addition of an option to select custom electrode subsets of a specified size and linear detrending for each response window. ERP responses windows were set for 0 to 800 ms post stimulus. The decimation frequency was set for downsampling to 20Hz. For classification, SWLDA was used with a maximum of 60 features, pEnter = .1, and pRemove = .15 [6]. SWLDA has long been reported to be effective for P300 classification with multichannel EEG [4].

In the Protocol 2 data, the amount of calibration data varied across participants due to difficulties in the online session with maintaining participant attention or in calibration. Some participants have fewer than 30 trials recorded, while some others have extra calibration data in the attempt to calibrate the BCI. Upon inspection, we found that the data was also more noisy than Protocol 1 data. We therefore applied 2 additional steps for preprocessing: 1) we chose a calibration file combination so that the total number of trials is as close to 30 as possible, and 2) we first tried to calibrate without spatial filtering, and if neither the default nor custom subset could calibrate, we applied a common average reference (CAR) filter.

### 2.5. Electrode Selection

For the electrode subset selection, we implemented the forward selection method proposed by McCann et al 2015 [9], which was shown to be as effective as exhaustive search when generalized to evaluation data. The forward selection algorithm added electrodes one-by-one that would locally maximize accuracy until the desired subset size was reached. Initially, all available electrodes were in the “available” set and the “chosen” set was empty. Electrodes were moved to the “chosen” set one at a time, based on the SWLDA calibration accuracies for the addition of one more electrode. Thus, each iteration obtained the calibration accuracies of all candidate subsets, each composed of all electrodes in the current “chosen” set plus one electrode in the “available” set. The candidate subset with the highest accuracy would be the new “chosen” set, and the corresponding electrode from the “available” set would be excluded from the new “available” set. Note that, although 8 electrodes are chosen for each custom subset, it is possible that some of the chosen electrodes do not contribute to the weight for classification. The electrodes that are used to generate weight is called “transmitted electrodes”.

### 2.6. Calibration

Calibration was done separately on data from each participant in both protocols 1 and 2. For consistency in data size across the two protocols, thirty trials from Protocol 1 were used. Calibration accuracy for the custom electrode subset and the default subset was compared. In case of a tie between the custom and default subset at 10 flashes, we decreased the number of flashes from 10 and report the accuracy at the first number of flashes when the tie was broken.

### 2.7. Testing

The larger amounts of data available for Protocol 1 enabled additional investigation of the ability of calibration data to generalize to testing data. For this investigation, 20 trials were selected randomly and held out for testing while the remaining forty trials were used for calibration. Electrode selection using the 40 calibration trials was done with an iterative process. The intention was to limit the effect of potential outlier trials. We treated electrode subset as a hyper-parameter, and applied a leave-one-out validation pipeline over the calibration data to fit the hyper-parameter. Specifically, for each participant, let EEG cap electrode size = S, calibration data = N selections with a target custom electrode subset size = M. One selection was held out at a time, and the remaining (N-1) selections were fitted with SWLDA with the forward algorithm for M electrodes. A total of N electrode subsets were obtained with the process. Then, each electrode received a score based on the order (k) in which it was added to the subset. Specifically, we found the top M highest scoring electrodes calculated by 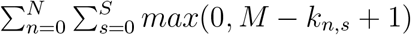 as the custom electrode subset for the participant.

This produced an electrode subset of size 8, which was then used to obtain weights for classification on the entire 40 selections of the calibration data. The weights were then applied to the held-out 20 selections to determine testing accuracy. Both calibration and testing accuracy were reported for participants in Protocol 1 who had complete data (9 participants with mild CP, 10 TD).

### 2.8. Statistical Analysis

Statistical analysis to assess the group effects and individual effects of the custom electrode subset was done in R version 4.0.4 [33].

#### 2.8.1. Group Effects

A generalized linear mixed-effects model (glmm) with poisson outcome was used to evaluate age-controlled group effects of electrode selection in both the calibration data and the testing data, with subset type (default or custom), group (typically developing controls, mild CP, severe CP), and age as fixed effects, and individual participant as a random effect.

A linear mixed-effects model (lmer) was used to assess whether the effect of electrode subset seen in calibration accuracy generalizes to accuracy on testing data. This analysis was done for the participants with complete data (10 people with mild CP, 10 typically developing controls). Difference in accuracy at 10 flashes (custom - default) was evaluated against the default subset accuracy, age, and the interaction between group and analysis paradigm (calibration vs testing).

#### 2.8.2. Individual Effects

We assessed the statistical significance of custom electrode subset on BCI accuracy at the individual level. For each participant, calibration data of an average of 30 trials from the default 8 electrodes was fitted to a binomial distribution using the maximum likelihood (MLE) method. It was assumed that each trial was independent and identically distributed (i.d.d.). Let the number of random trials = n, and the number of correct classifications using the default subset = k. We calculated the cumulative probability *P*(*X* ≥ *k*) from the custom 8 electrodes with respect to the fitted binomial distribution. This statistic translates as “using the custom subset, in n random trials, the probability of seeing greater or equal to k success”, which entails whether a custom subset improves BCI accuracy when compared to the default subset. The cumulative probability is equivalent to a one-sided p-value using the binomial distribution.

Distance between individualized subset and the default subset is quantified by a variant of the Earth Mover’s Distance (EMD) metric, detailed in Pele and Werman 2008 [34]. EMD represents the minimal distance between 2 sets of data points by mapping set A to set B. The EMD variant used in this paper adds a penalty term to cover the cases when the number of points is different between set A and B, since the number of transmitted electrodes can vary across participants. The implementation of the algorithm from https://github.com/wmayner/pyemd is used in this study.

## 3. Results

### 3.1. Calibration Accuracy

At the group level, between group comparison showed that custom electrode subset significantly improved BCI calibration accuracy for the severe CP group (p < 0.0001), but not for the mild CP group (p = 0.185) nor the controls (p = 0.676). For the severe CP group, the average accuracy using the custom subset improved by 28.6% (95% CI [13.4%, 46.1%]) compared to the default subset. Using the default subset, there was no significant difference in calibration accuracy between the mild CP group and the controls (p = 0.208), but the BCI accuracy was significantly worse in the severe CP group than the controls (p < 0.0001). Using the custom subset, there was no significant difference between the mild CP and the severe CP group (p = 0.522), but the BCI for the control group performed significantly better (p = 0.0005). Figure 4 shows changes in calibration accuracy by electrode subset (custom vs default) for individuals in each group. within group comparison shows that age significantly affected calibration accuracy (p = 0.020) (Figure 3).

**Figure 3:**
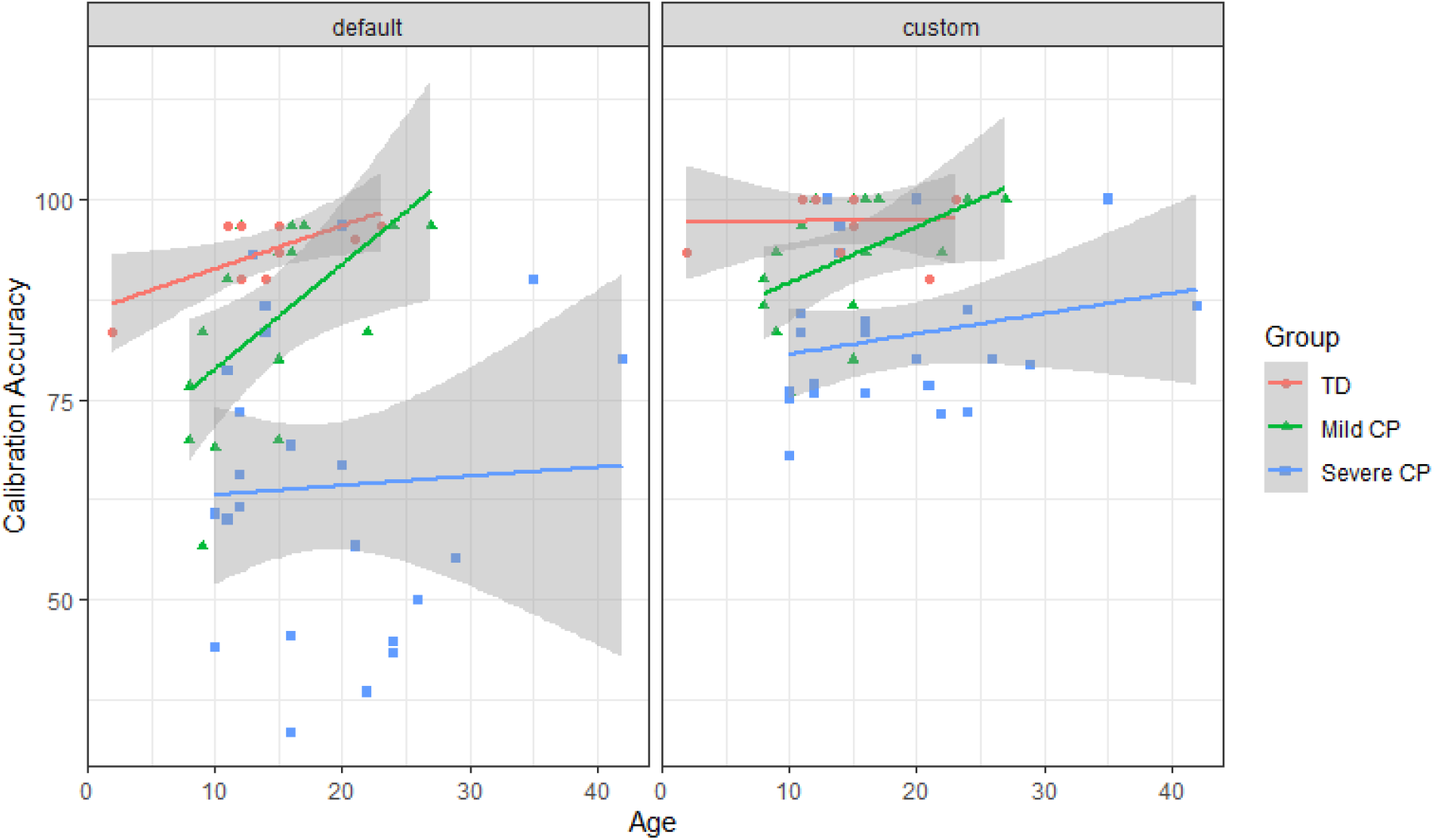
BCI calibration accuracy across age. Experimental data is plotted as scatters and a fitted line is plotted for each group for each subset type. The shaded area represents the 95% confidence interval.

**Figure 4:**
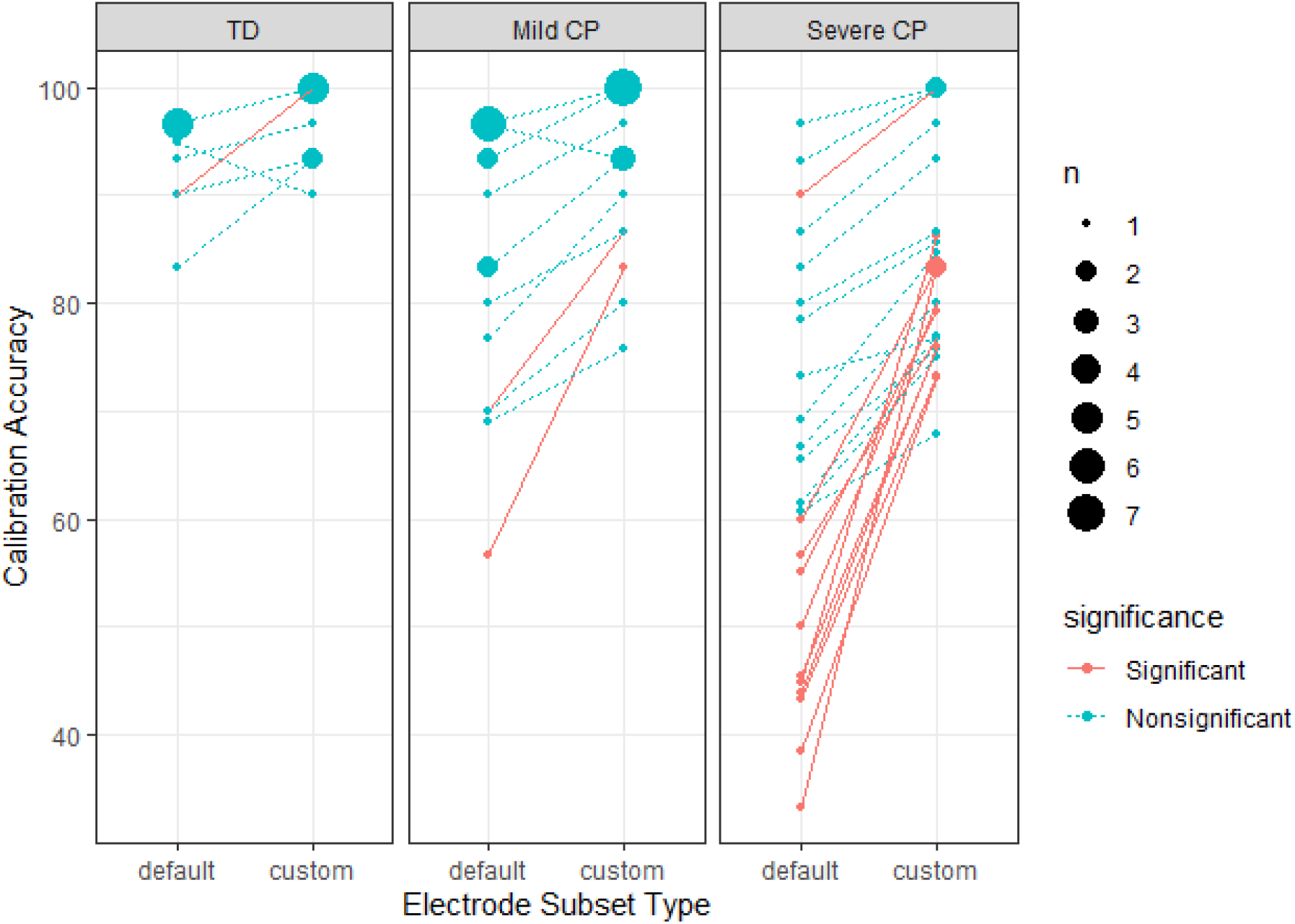
BCI calibration accuracy with different electrode subset types by participant group. The solid pink lines represent individuals with a significant improvement calculated as described in section 2.8.2. The size of the data points represents the number of participants with the same accuracy score. Note, that the magnitude of improvement is not a complete representation of the statistical significance.

**Figure 5:**
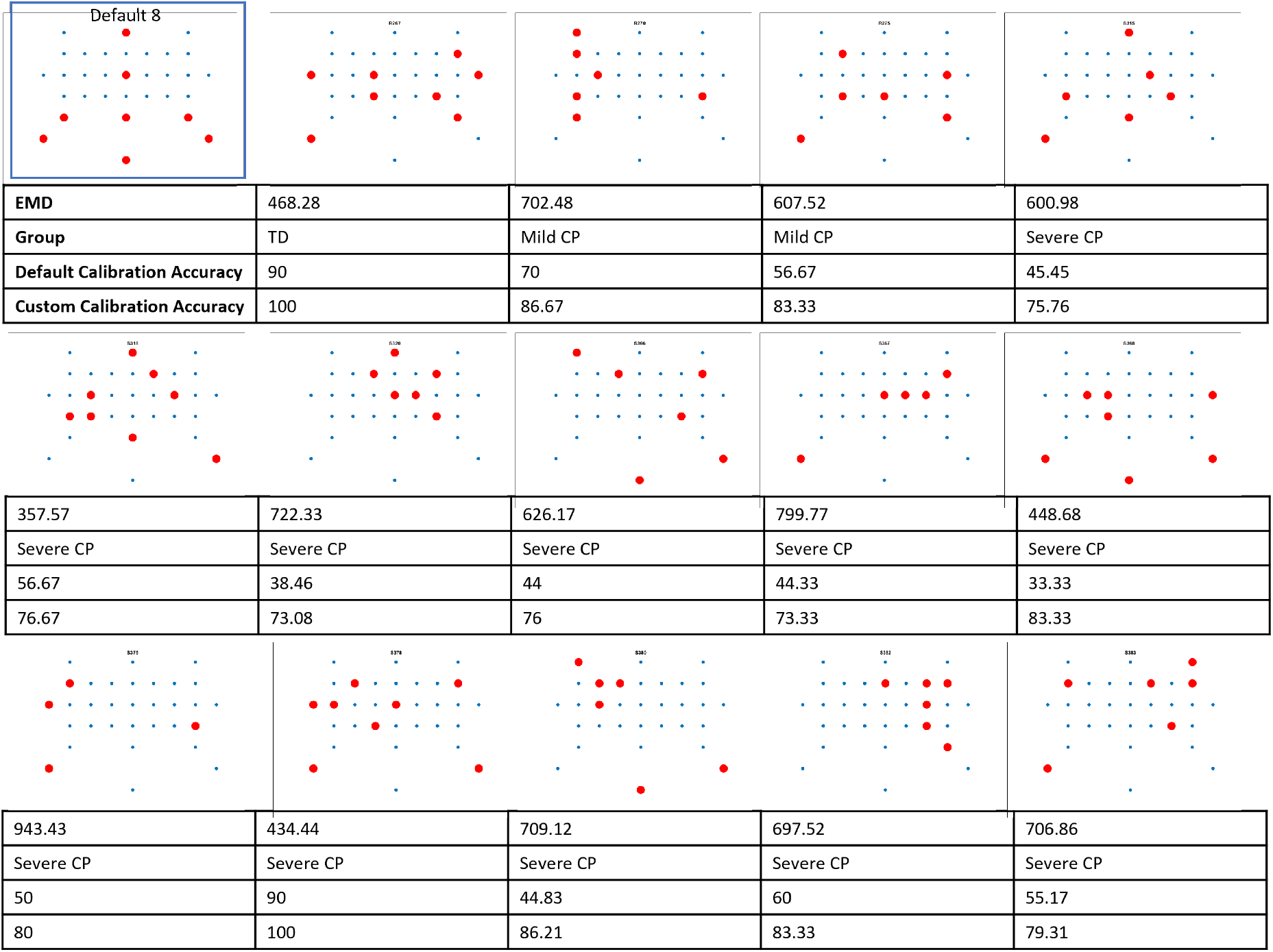
Custom subset locations, default and calibration accuracy for each participant with significant improvement. Together they show the degree of variability of custom electrode locations and the resulting changes in BCI performance.

For individual effects, electrode selection produces significant improvement in calibration accuracy for one control (out of 10 participants), two people with mild CP (out of 17 participants), and 11 people with severe CP (out of 24 participants) (see figure 4, figure 5). It is worth noting that the two people with mild CP who had significant improvement did not meet minimum calibration accuracies with the default subset during the data collection to proceed to testing.

### 3.2. Testing Accuracy

Comparing the default and custom subsets, neither the calibration accuracy with the default subset (p = 0.160) nor age (p= 0.488) significantly predicted the change in BCI accuracy from calibration to testing. For the controls, there was a significant reduction (—9.58%, 95% CI [—13.33%, —5.84%]) in the effect of the subset during testing compared to calibration (p < 0.0001). There was no evidence that the effect of electrode subset on testing accuracy was different from its effect on calibration accuracy in the mild CP group (p = 0.556).

We evaluated significance of group effects of electrode selection on testing accuracy. Comparisons were done using the number of flashes where ties are broken. Neither the mild CP (p = 0.694) nor the control group (p = 0.495) showed significant difference in testing accuracy between using the custom and default subset (figure 6).

**Figure 6:**
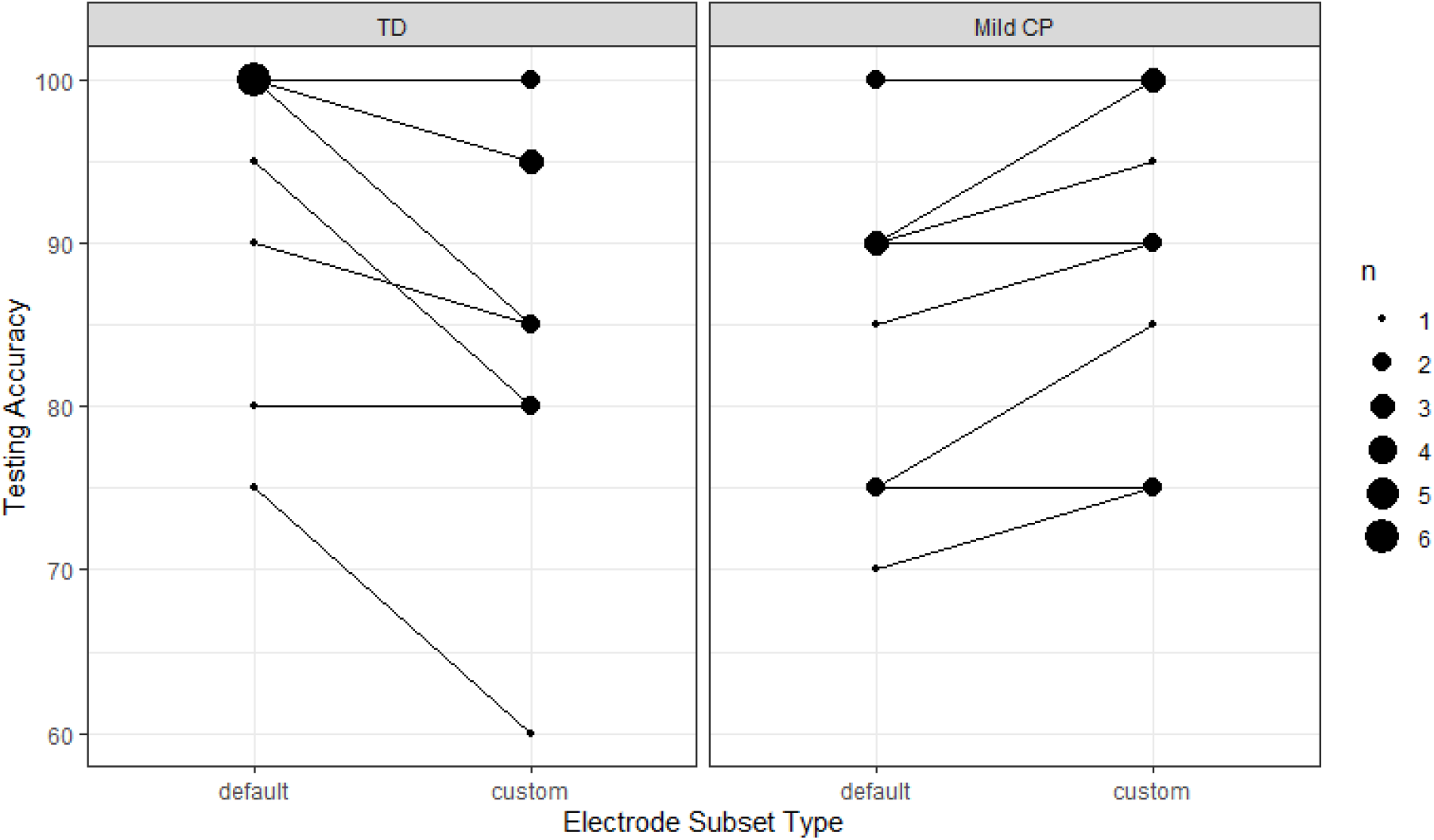
BCI testing accuracy with different electrode subset types. No significant improvement is found at the individual nor the group level. The size of the data points represents the number of participants with the same accuracy score.

### 3.3. Number of Transmitted (Useful) Electrodes

Differences in calibration accuracy as the size of custom electrode subset increases were investigated (figure 8a). As found by McCann et al. [9], most participants had an asymptotic increase in accuracy as electrode subset sizes increase. People with severe CP needed a bigger electrode subset size to achieve at least 95% of their own maximum calibration accuracy at 8 electrodes (figure 8b).

**Figure 7:**
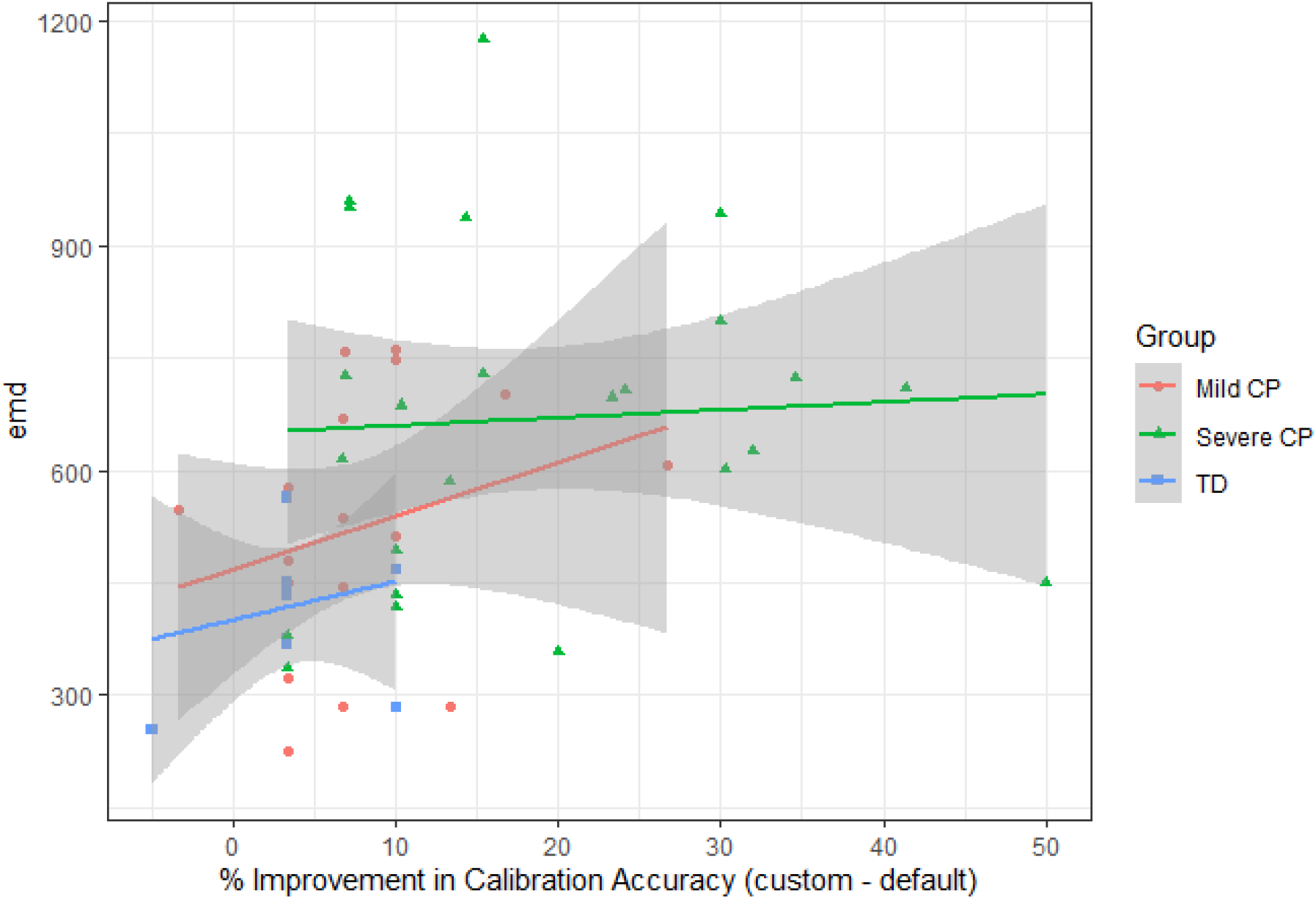
EMD with respect to improvement in calibratino accuracy by using an individualized electrode subset by group.

**Figure 8:**
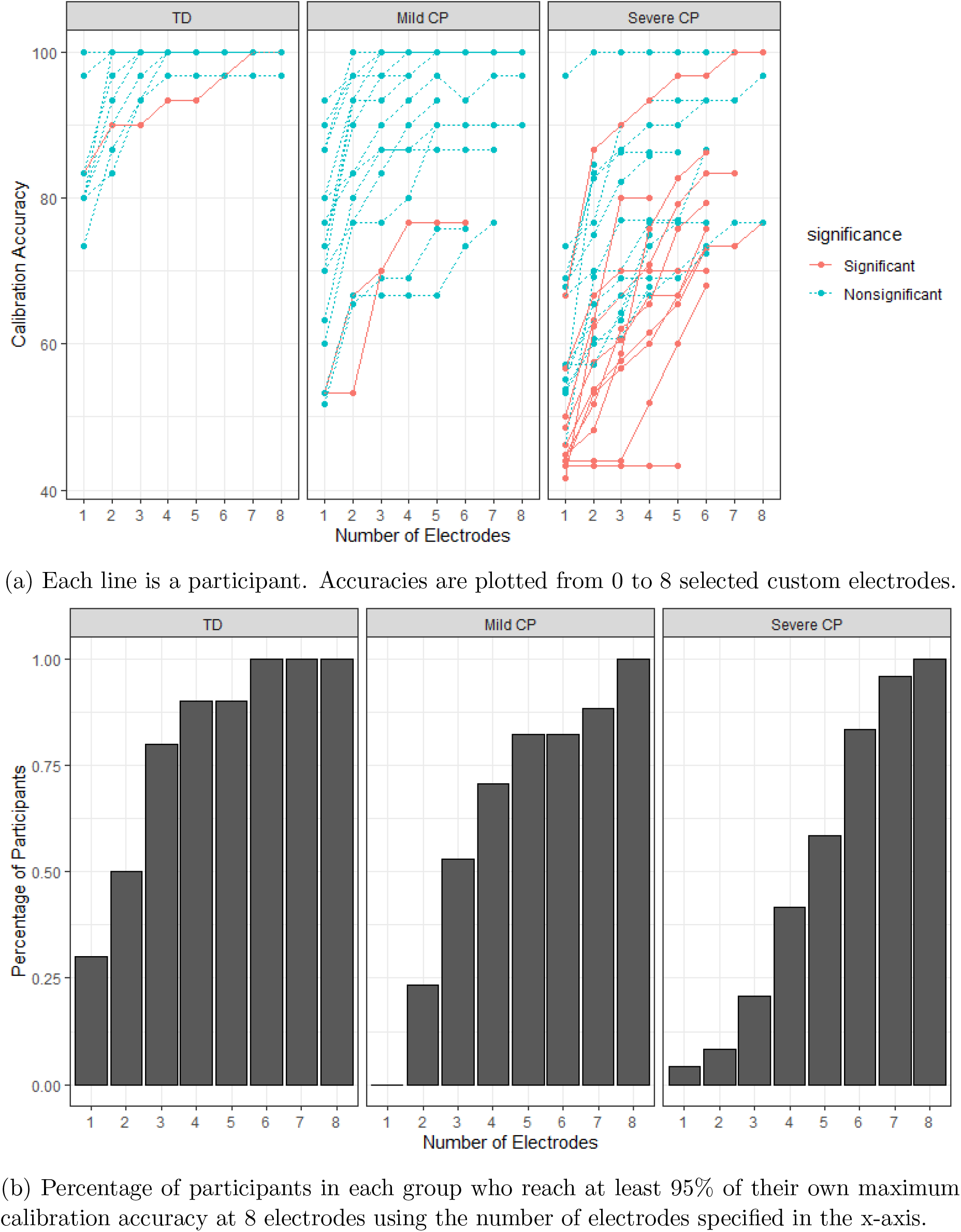
Calibration accuracy for custom electrode subset sizes.

Overall, limiting the maximum electrode subset size to 8 is efficient. Of the 14 people who saw improved performance for the custom electrodes, 11 of them did not use all 8 custom electrodes. While the remaining 3 might have gotten some benefit from additional electrodes, for most people a maximum of 8 electrodes is sufficient.

### 3.4. Distance between Individualized Custom Subsets and Default Subsets

The average EMD between custom and defaut subsets is 668.17 in the severe CP group, 524.33 in the mild CP group, and 419.75 in the typically developing group.

## 4. Discussion

The findings indicate that an individualized electrode subset can be essential for BCI calibration for people with severe CP, and an individualized electrode subset can benefit people with mild CP. Figure 3 shows improvement from the custom electrode subset is not limited to a particular age. However, the default electrode subset is adequate or even preferred for typically developed individuals.

### 4.1. Selected Electrodes and ERPs

Fig. 5 shows the custom electrode locations for each participant whose BCI calibration accuracy improved significantly with electrode selection. We notice that for some participants with CP, the selected electrodes are heavily lateralized to one side of the brain. There is a trend for the EMD values to be greater for participants with greater improvement using the custom subset as shown in 7, suggesting that the useful electrodes for some participants with CP are not covered in the default subset.

The etiology and pathophysiology of CP may explain and greater EMD values in participants with CP and the benefit gained from a custom electrode subset. CP stems from an early brain lesion and can result in neurophysiological differences including differences in representation of functions [35, 36]. For example, people with spastic CP often show evidence of periventricular leukomalacia (PVL) [37]. PVL has been associated with altered connectivity including reduced posterior pathways to parietal and occipital regions. Therefore, electrode selection may identify better custom locations to record informative EEG to accommodate those differences. Our result suggests that there are electrodes that are not in the default subset that are in fact informative for some participants with CP, and our electrode selection algorithm is able to identify those electrodes.

Fig. Appendix A.1 shows the average EEG for each electrode for a representative participant from each group. We did not aggregate ERPs at the group level because we expect individual differences in the ERP in people with CP. The red boxes (Fig. Appendix A.1) indicate the selected electrodes, showing that electrode selection successfully selects electrodes that better discriminate between targets and non-targets. From visual inspection, we can see that the data from the severe CP group is more noisy than the other two groups, probably due to movements and wandering attention as noted during the experiment. The ERPs also show a more restricted spatial distribution for the participant with severe CP, indicating a greater sensitivity to individual electrode selection. Electrode locations may be somewhat interchangeable for the control and mild CP groups due to the widespread nature of the ERPs.

### 4.2. Number of Useful Electrodes

McCann et al. [9] found minimal improvements in BCI accuracy for electrode subsets larger than 4 electrodes. Our results with the typically developing group support that finding, since at subset size 4, 9 out of 10 control participants were within 95% of their maximum accuracy and all had reached that threshold by subset size 6. However, this conclusion cannot generalize to people with CP, as Fig. 8b shows that over 50% of the participants with severe CP benefit from using more than 4 electrodes and for some a subset of 8 electrodes is more effective than 7.

### 4.3. Electrode Selection for Typically Developing Individuals

Colwell et al. [6] previously reported significant improvement for typically developing individuals with sequential forward search; however, we were not able to replicate this result. The research focus in Colwell et al. concerned whether an electrode set could improve BCI performance for all participants, while our goal was to evaluate the need for a custom individualized electrode subset for each participant. The major differences in methods between our study and Colwell et al., are: 1) experimental paradigm, 2) the electrode locations on the EEG cap, and 3) the age of participants.

The experimental paradigm is different both in the number of selections and the number of flashes. Colwell et al. uses the P300 Speller 9×8 Checkerboard paradigm with 38 character selections for calibration, while our study used a 4-question multiple-choice paradigm with individual flashes.

Our electrode set was designed to have more coverage in the motor cortex to allow for more flexibility for people with spastic CP. The electrode locations available in Colwell et al. contain more posterior electrodes compared to our electrode cap, and many of the most useful electrodes in Colwell et al. for their typically developing participants are posterior electrodes. We speculate that, because all of the posterior electrodes in our study were already in the default subset, there was no additional posterior electrodes for the control group to leverage to further improve BCI accuracy.

Age is not reported in Colwell et al, but we can infer from ‘‘received course credit in return for participation in this study” that participants are college-age young adults. Our typically developing participants were younger with a wider range of ages (14.7 ± 4.3 years). The conflicting results suggest that effectiveness of electrode selection could be task and/or search space and/or age dependent, and caution is needed when applying electrode selection methods to a different population for a different experiment on a different system.

Furthermore, a motor-imagery BCI study [38] showed that typically developing individuals do not benefit from electrode selection in online studies, though they did find improvement in an offline study. Meng et al. suggested that non-task related modulation (captured by selected electrodes that are located outside the targeted cortices) could result in better BCI performance in a single session, but may not be beneficial for skill consolidation in long-term BCI use [38]. These inconsistent results are a reminder that there is no clear physiological explanation for benefit from electrode selection for typically developed individuals.

### 4.4. Limitations

Electrode selection was not available online during the two protocols. Data from many participants with severe CP were noisy and thus additional preprocessing was applied to all participants with severe CP. Brain imaging scans were not available for our participants, so we could not directly compare selected electrodes or ERP locations to neuro-anatomy.

## 5. Conclusion

The study shows for the first time that people with severe CP can benefit significantly from electrode subset selection. This study’s results with typically developing individuals conflict with reports from the literature, suggesting that the effectiveness of electrode selection may be subject to the experimental condition, age and the available electrodes.

## 6. Acknowledgement

The content is solely the responsibility of the authors and does not necessarily represent the official views of the funding sources.

Research supported by 1) the Mildred E. Swanson Foundation and by the National Center for Advancing Translational Sciences of the National Institutes of Health under Award Number UL1TR000433. The content is solely the responsibility of the authors and does not necessarily represent the official views of the Mildred E. Swanson Foundation or the National Institutes of Health; 2) Michigan Institute for Clinical & Health Research grant support (CTSA: UL1TR002240); and 3) Cerebral Palsy Alliance Research Foundation.

The authors would like to thank Corey Powell, consultant at Consulting for Statistics, Computing and Analytics Research (CSCAR) at the University of Michigan, for his support on statistical analysis in this manuscript.

Study data were collected and managed using REDCap electronic data capture tools hosted at the University of Michigan. [39, 40]REDCap (Research Electronic Data Capture) is a secure, web-based software platform designed to support data capture for research studies, providing 1) an intuitive interface for validated data capture; 2) audit trails for tracking data manipulation and export procedures; 3) automated export procedures for seamless data downloads to common statistical packages; and 4) procedures for data integration and interoperability with external sources.

## Appendix A

**Figure Appendix A.1:**
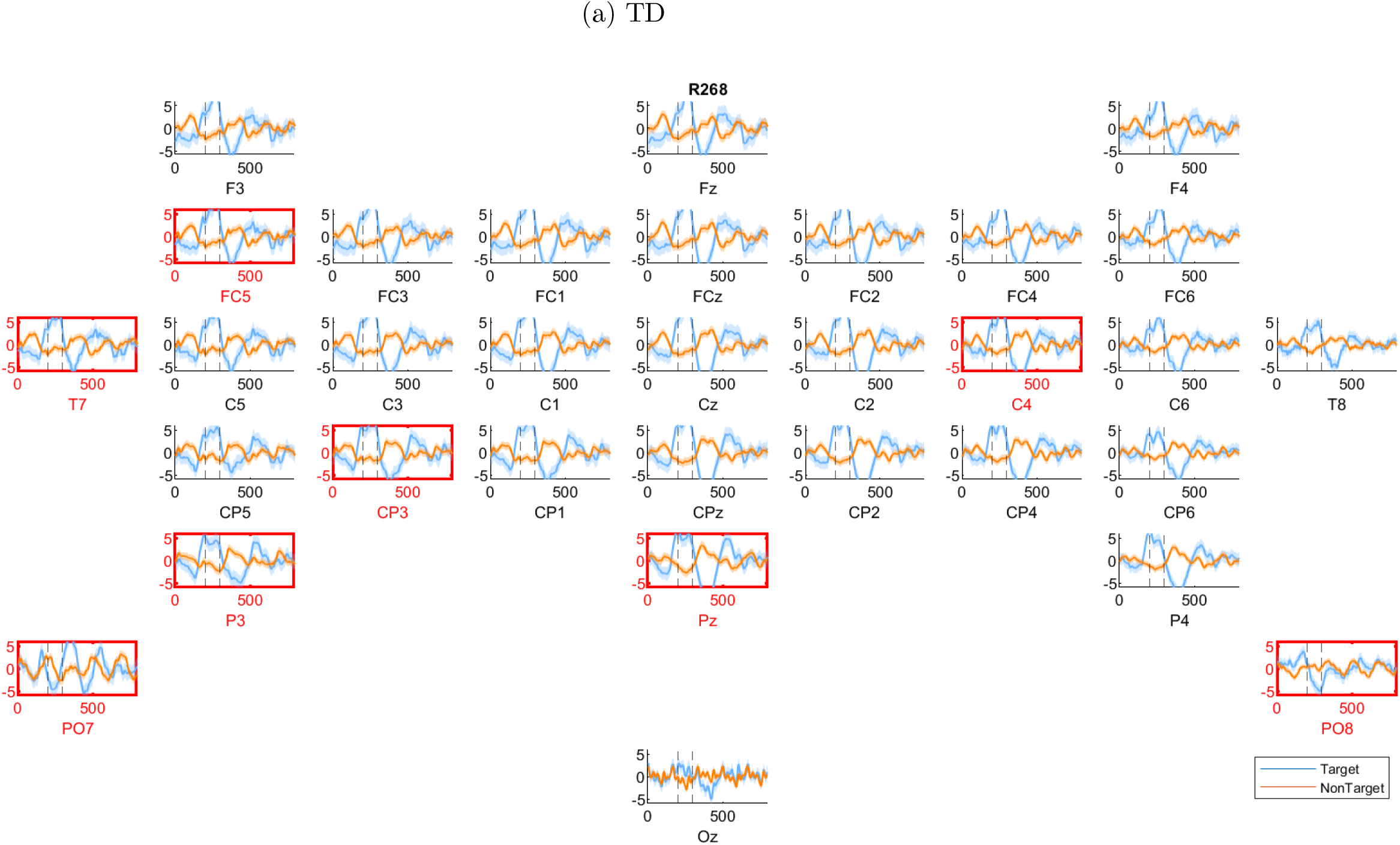
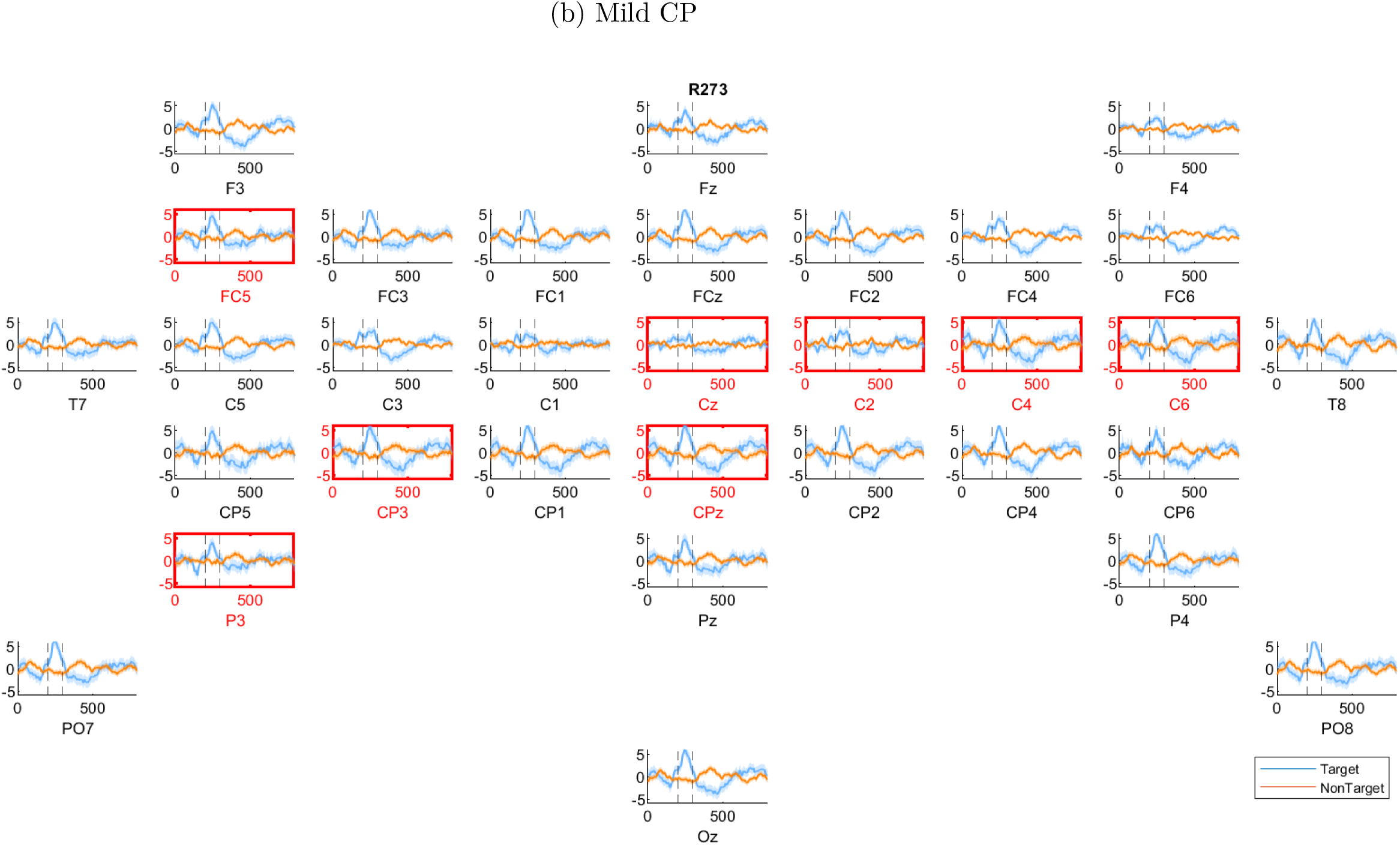
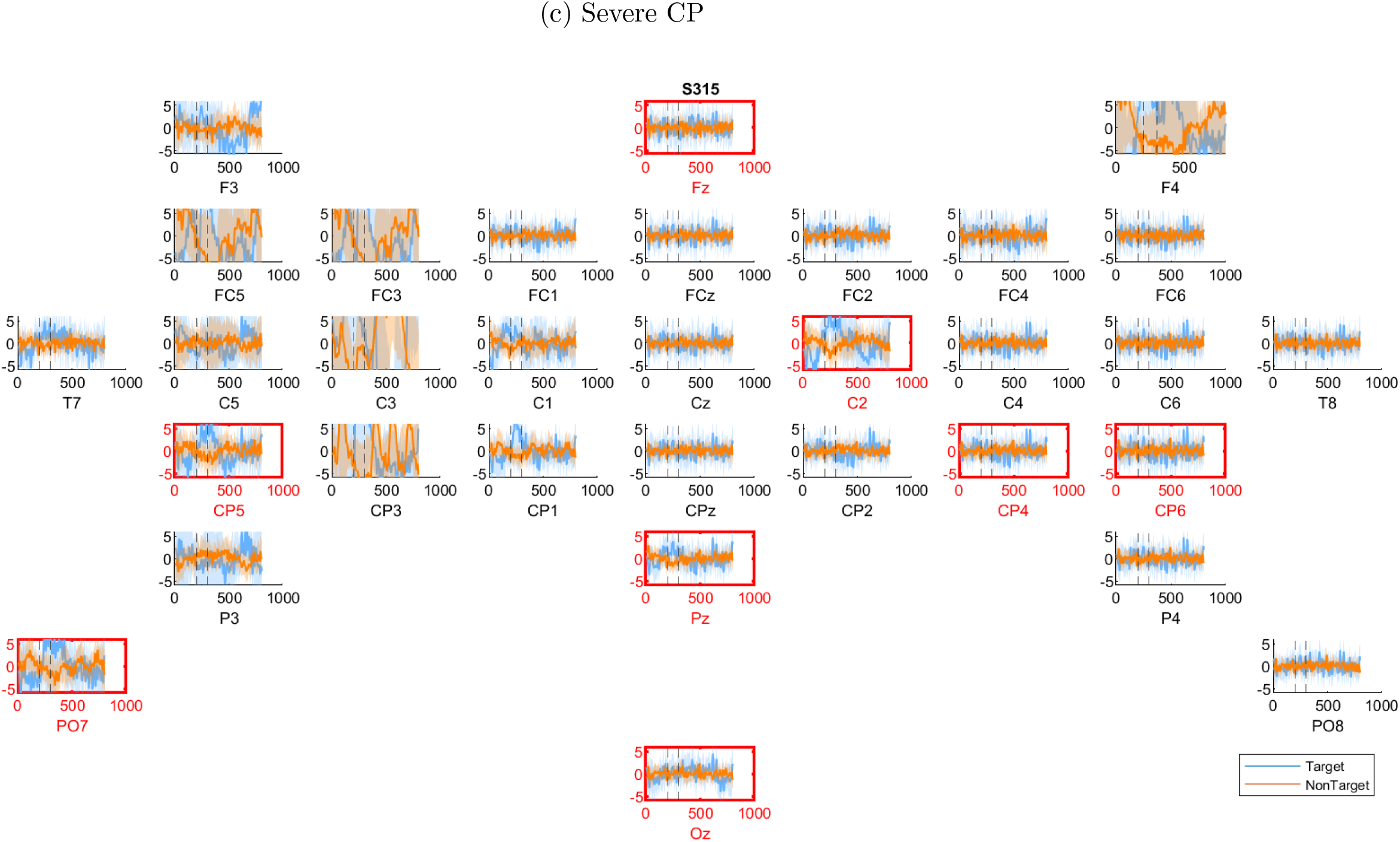
Average target (blue) and non-target (orange) EEG of 30 trials for 0 to 800ms post stimulus. The 30 trials produce 300 target responses and 900 non-target responses. The shaded region represents the 95% confidence interval. The dotted reference lines mark t = 200 ms and t = 300 ms. Red boxes indicate the selected custom electrodes for the participant, (a) Example from the control group; (b) Example from the mild CP group; (c) Example from the severe CP group.

